# Gene coordination patterns across 8,314 tumors reveal a spectral point of no return in cancer progression

**DOI:** 10.64898/2026.04.30.722092

**Authors:** John Mayfield

## Abstract

Genes operate through coordinated patterns that preserve tissue self-identity, regulate immunity, and control growth. We hypothesize that this coordination undergoes a structured collapse during cancer progression and that there is a shared point of no return across multiple cancer types that may help predict patient outcomes. We calculated gene–gene coordination across 8,314 tumors spanning 32 cancer types by measuring how similarly each pair of 3,000 genes behaves across the patient population. We find that 71% of all coordination is captured by a single pattern, with three secondary patterns that predict survival in 11 of 30 cancer types including patterns involving metabolic dedifferentiation (cytochrome P450 metabolism vs. epithelial differentiation), immune polarization (adaptive vs. innate immunity), and tissue self-identity. Specifically, this method stratifies prognosis in glioblastoma (C-index 0.845, *p* = 1.8 × 10^−6^), prostate adenocarcinoma (0.800), clear cell renal carcinoma (0.649), and eight additional cancers (*p* = 6.9 × 10^−32^). Beyond this threshold, tumors of all types share uniformly poor survival, defining a universal molecular point of no return and a potential window for early intervention before coordination collapse becomes irreversible.

## Introduction

The molecular events that drive cancer progression (somatic mutations, copy number alterations, epigenetic reprogramming) have been catalogued at extraordinary resolution across tens of thousands of tumors. Yet a basic question remains unanswered: at what point does the accumulation of these events become irreversible? Clinicians recognize this transition intuitively. A low-grade glioma that transforms to glioblastoma, a localized prostate cancer that acquires metastatic potential, an indolent renal mass that begins aggressive growth. In each case, there is a qualitative shift from a state of disordered-but-contained biology to a state of coordinated malignant behavior. We lack a quantitative framework for detecting this shift directly from molecular data.

The difficulty is not a shortage of data but a mismatch between the analytical tools available and the biological question being asked. Principal component analysis (PCA) and related methods identify axes of maximal variance across patients, answering the question “which patients differ most from each other?” Clustering algorithms partition patients into discrete subtypes, answering “which patients are similar?” Neither addresses the question that matters for understanding cancer progression: which genes coordinate with each other, and how does that coordination change as a tumor advances?

This distinction is not semantic. Variance and coordination are mathematically different quantities. PCA decomposes the sample-by-sample covariance matrix, whose eigenvectors are patient groupings. Gene–gene coordination requires decomposing the feature-by-feature similarity matrix, whose eigenvectors are groups of genes that move together. A gene that is highly variable across patients may or may not participate in a coordination program; a gene with modest variance can be a central node in a tightly coordinated module. The coordination structure is invisible to PCA.

Here we propose that cancer progression corresponds to a reorganization of the gene–gene coordination landscape, a transition from a distributed architecture, in which many independent programs coexist, to a monopolar architecture dominated by a single coordination mode. This transition is directly analogous to a phase transition in statistical mechanics, where a system moves from a disordered state with many accessible configurations to an ordered state dominated by one. We formalize this analogy by constructing a Hamiltonian, the operator that encodes all pairwise interactions in a physical system, from the gene–gene similarity kernel, and we decompose it into its eigenmodes.

The eigenmodes of this Hamiltonian are not variance axes. They are coordination programs: sets of genes whose expression levels rise and fall together across the 8,314-patient pan-cancer population. Their eigenvalues measure the strength of each coordination program, specifically how tightly the constituent genes are coupled. The spectral concentration *c*, defined as the fraction of total coordination carried by the dominant eigenmode, quantifies whether the system is in a distributed (low *c*) or monopolar (high *c*) regime. And the critical temperature *T* ^*^, computed from the spectral gap, identifies the boundary between these regimes.

We test three predictions of this framework. First, if cancer progression involves coordination collapse, then the pan-cancer landscape should show high spectral concentration—a single program should dominate. Second, the secondary coordination modes (those not captured by the dominant program) should have interpretable biological identities recoverable by gene set enrichment analysis. Third, a patient’s position along these coordination modes, their spectral malignancy coordinate *η*, should predict overall survival, with the direction of the effect depending on which coordination program dominates in each cancer type.

## Results

### The pan-cancer coordination landscape is monopolar

We computed the gene–gene coordination kernel for 8,314 tumors across 32 cancer types using 3,000 variance-filtered mRNA expression features from the MLOmics pan-cancer compendium. For each pair of genes *i* and *j*, we measured their coordination across patients using a radial basis function (RBF) kernel:

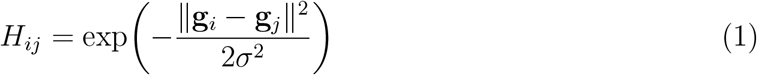

where g_*i*_ ∈ ℝ^8314^ is the expression profile of gene *i* across all patients (z-scored) and *σ* is set by the median heuristic (*σ* = 126.6). The resulting 3,000 × 3,000 Hamiltonian *H* is a symmetric positive semi-definite matrix whose eigenmodes represent coordination programs.

Eigendecomposition of *H* revealed a striking monopolar architecture (Fig. 1A,C). The dominant eigenvalue (*λ*_1_ = 1,844.8) was 12.6-fold larger than the second (*λ*_2_ = 146.9), yielding a spectral concentration of *c* = 0.71 and a spectral gap of Δ*λ* = 1,697.8. This means that 71% of all gene-gene coordination across the entire pan-cancer transcriptome is captured by a single program. The coordination landscape is not distributed across many independent modules; it is dominated by one.

**Figure 1.**
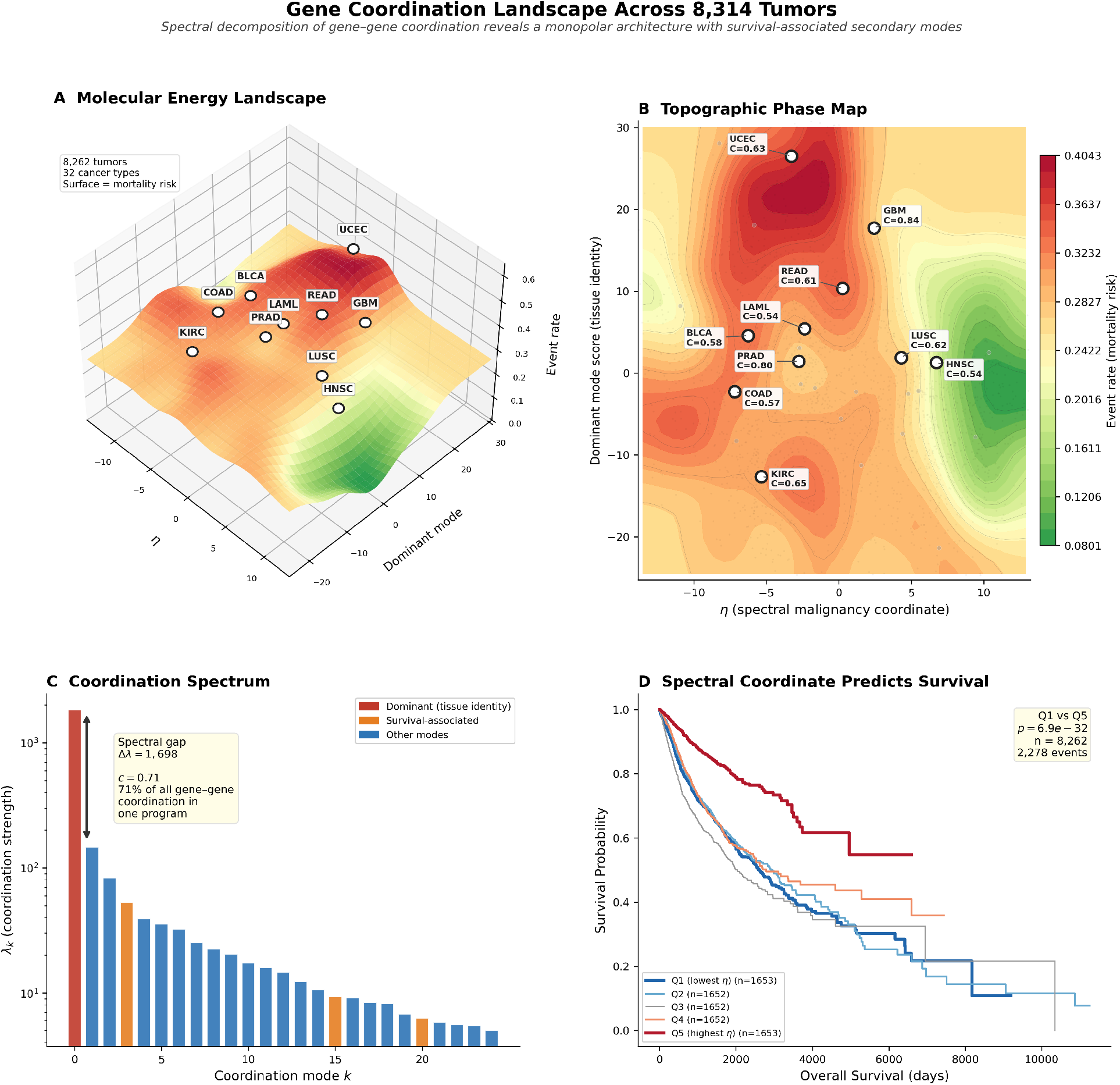
Gene coordination landscape across 8,314 tumors. **(A)** Three-dimensional molecular energy landscape. The surface represents binned event rate (mortality risk) as a function of the spectral malignancy coordinate *η* (*x*-axis) and dominant coordination mode score (*y*-axis) across 8,262 patients with survival data. Cancer centroids for the 11 significant types (*p <* 0.05) are marked with labeled red circles; non-significant types are shown as gray crosses. The ridge of elevated event rate identifies the high-risk region of coordination space. **(B)** Topographic phase map (top-down projection of the energy landscape). White circles mark significant cancer centroids with C-index annotations. Surface contours indicate event rate isolines. The spatial separation of cancer types reflects their distinct positions in the gene coordination landscape. **(C)** Coordination spectrum showing eigenvalues on a log scale. The dominant eigenmode (*λ*_1_ = 1,845, red) captures 71% of all inter-gene coordination (*c* = 0.71), with a spectral gap Δ*λ* = 1,698 to the second mode. Orange bars mark the three survival-associated coordination modes (modes 3, 15, 20). **(D)** Pan-cancer Kaplan–Meier survival curves stratified by *η* quintile across all 8,262 patients with survival data. Q1 vs. Q5 log-rank *p* = 6.9 *×* 10^*−*32^; *n* = 8,262; 2,2784events.

This monopolar architecture has a direct physical interpretation. In the language of statistical mechanics, it corresponds to a system deep in its ordered phase, well below the critical temperature *T* ^*^ = 1*/*Δ*λ*. The partition function 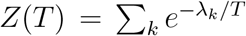 concentrates overwhelmingly on the ground state. The biological implication is that the pan-cancer transcriptome, when viewed through the lens of gene coordination rather than gene variance, operates in a highly ordered regime where most genes participate in a single dominant program. The 3D molecular energy landscape and its 2D topographic projection (Fig. 1A,B) reveal that cancer types with the strongest survival associations occupy distinct regions of this coordination space, with event rate varying systematically across the spectral coordinate *η* and the dominant mode score.

### Secondary coordination modes have distinct biological identities

While the dominant mode captures tissue-of-origin coordination, the survival-relevant biology resides in the secondary modes—the coordination programs that operate orthogonally to the dominant one. Screening the top 30 modes for survival association (by concordance index against overall survival across all 8,262 patients with follow-up) identified three modes with C-index *>* 0.58: modes 3, 15, and 20.

Gene set enrichment analysis (GSEA) on the gene loadings of each mode revealed biologically coherent coordination programs (Fig. 2; Supplementary Tables 1–3):

**Figure 2.**
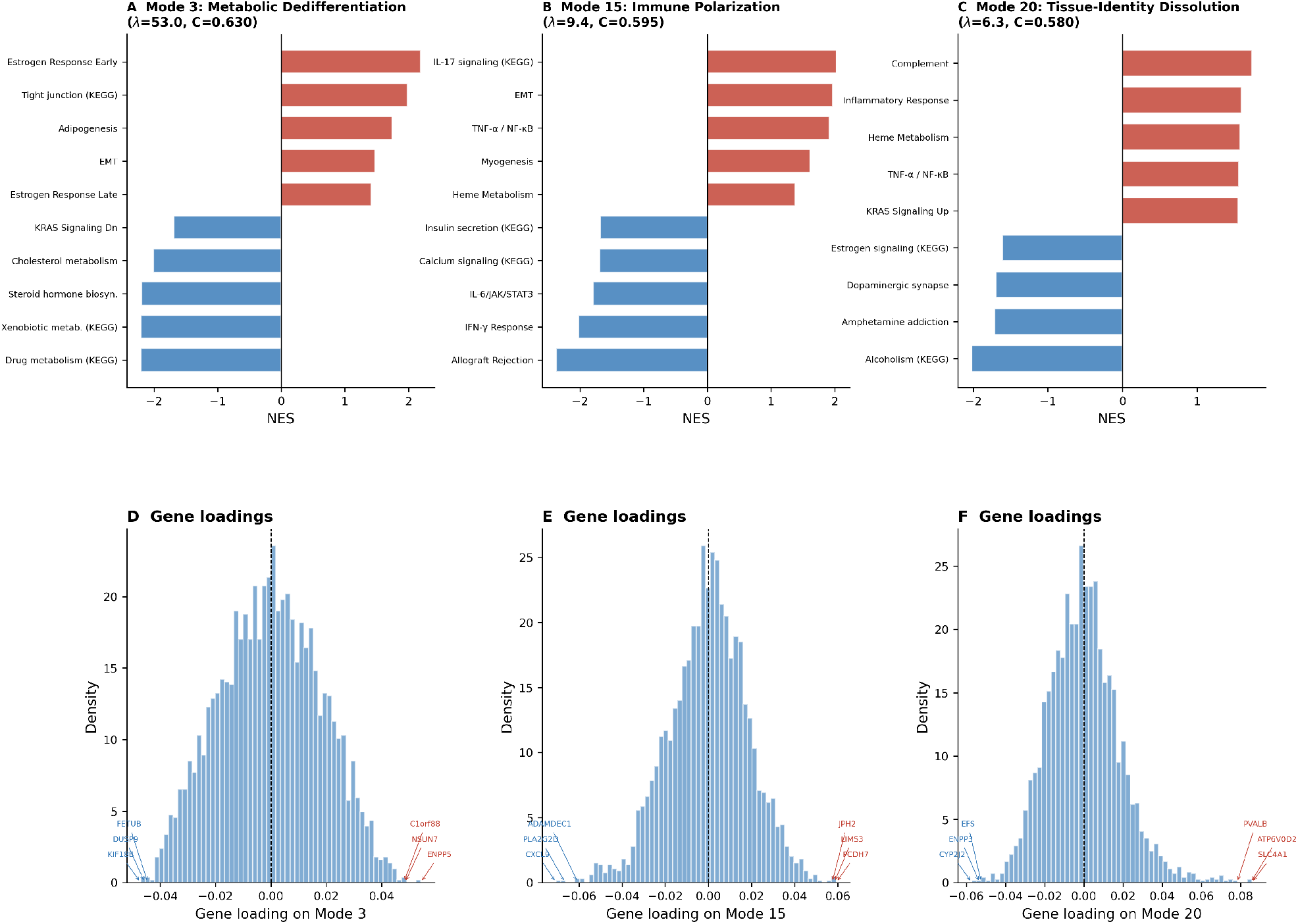
Three survival-associated coordination modes represent metabolic dedifferentiation, immune polarization, and tissue-identity dissolution. **(A–C)** Gene set enrichment analysis (pre-ranked GSEA, MSigDB Hallmark 2020 and KEGG 2021 Human) on gene loadings of the three coordination modes with highest survival concordance index. Red bars: positive normalized enrichment score (NES); blue bars: negative NES. All terms shown at FDR *q <* 0.25. **Mode 3** (*λ* = 53.0, C-index = 0.630) separates epithelial differentiation programs (estrogen response, NES = +2.19; tight junctions, NES = +1.98) from metabolic reprogramming (cytochrome P450 drug metabolism, NES = *−*2.21; xenobiotic metabolism, NES = *−* 2.21; steroid hormone biosynthesis, NES = *−* 2.19). **Mode 15** (*λ* = 9.4, C-index = 0.595) separates adaptive immunity (allograft rejection, NES = *−*2.37; IFN-*γ* response, NES = −2.02) from innate inflammatory signaling (IL-17, NES = +2.02; TNF-*α*/NF-*κ*B, NES = +1.91; EMT, NES = +1.97). **Mode 20** (*λ* = 6.3, C-index = 0.580) captures complement and inflammatory programs versus neuroendocrine signaling. **(D– F)** Distribution of gene loadings on each coordination mode, with top-contributing genes annotated. Mode 3 loadings show mitotic kinases (KIF18B, KIF14, AURKB) on the negative pole and tight junction/epithelial markers (CLDN3, C2orf40) on the positive pole.

#### Mode 3: metabolic dedifferentiation (*λ* = 53.0, C-**index** = 0.630)

The positive pole (associated with better survival) enriched for estrogen response (NES =+2.19, FDR *<* 0.001), tight junctions (NES = +1.98, FDR = 0.038), and adipogenesis (NES = +1.74, FDR = 0.023)—programs of epithelial differentiation and tissue-specific metabolic identity. The negative pole enriched for cytochrome P450 drug metabolism (NES = −2.21, FDR *<* 0.001), xenobiotic metabolism (NES = −2.21, FDR *<* 0.001), steroid hormone biosynthesis (NES = −2.19, FDR *<* 0.001), and cholesterol metabolism (NES = −2.01, FDR = 0.003). The top gene loadings on the positive pole included tight junction components (CLDN3) and epithelial markers (C2orf40, TMC4); the negative pole included mitotic kinases (KIF18B, KIF14, AURKB) alongside metabolic enzymes. This mode captures a coordination axis between tissue differentiation and metabolic reprogramming: tumors in which these programs are coordinated toward the differentiation pole retain their tissue identity and have better outcomes, while those coordinated toward the metabolic/proliferative pole have undergone dedifferentiation.

#### Mode 15: immune polarization (*λ* = 9.4, C-index = 0.595)

The negative pole enriched for interferon-*γ* response (NES = −2.02, FDR = 0.002) and allograft rejection (NES = −2.37, FDR *<* 0.001)—hallmarks of cytotoxic adaptive immunity. Top gene loadings on this pole included CXCL9, CD8B, CD79A, and ADAMDEC1. The positive pole enriched for epithelial–mesenchymal transition (NES = +1.97, FDR = 0.003), TNF-*α*/NF-*κ*B signaling (NES = +1.91, FDR = 0.004), and IL-17 signaling (NES = +2.02, FDR = 0.012), with top gene loadings including FOSL1, EREG, and TREM1. This mode separates tumors with coordinated adaptive immune microenvironments from those with coordinated innate inflammatory signaling—the immune polarization axis that determines checkpoint inhibitor eligibility.

#### Mode 20: tissue-identity dissolution (*λ* = 6.3, C-index = 0.580)

The positive pole enriched for complement (NES = +1.73), inflammatory response (NES = +1.59), and IL-6/JAK/STAT3 (NES = +1.51), with gene loadings dominated by renal-specific transporters (SLC4A1, ATP6V0A4, CLCNKB) and tissue-specific enzymes (CYP11A1, KLK1). This mode captures whether a tumor retains coordinated expression of its tissue-of-origin functional genes or has dissolved that coordination, explaining its particularly strong performance in kidney clear cell carcinoma (KIRC, C-index = 0.649).

### The spectral malignancy coordinate stratifies prognosis across cancer types

We defined the spectral malignancy coordinate *η*_*i*_ for each patient *i* as the mean projection onto the three survival-associated coordination modes, normalized by spectral concentration:

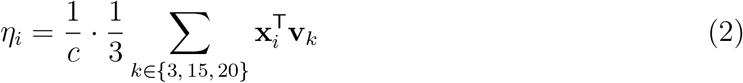

where x_*i*_ ∈ ℝ^3000^ is the z-scored expression profile of patient *i*, v_*k*_ is the *k*-th eigenvector of *H* (a coordination pattern in gene space), and *c* is the spectral concentration. The division by *c* normalizes for the degree of monopolar dominance, placing *η* on a scale that is comparable across datasets with different coordination architectures.

The coordinate *η* stratified overall survival in 11 of 30 evaluable cancer types at *p <* 0.05 (Table 1; Fig. 3). The strongest associations were: glioblastoma (GBM, C-index = 0.845, *p* = 1.8 × 10^−6^), prostate adenocarcinoma (PRAD, C-index = 0.800, *p* = 0.009), kidney clear cell carcinoma (KIRC, C-index = 0.649, *p* = 7.4 × 10^−5^), uterine corpus endometrial carcinoma (UCEC, C-index = 0.634, *p* = 0.009), lung squamous cell carcinoma (LUSC, C-index = 0.618, *p* = 0.023), rectal adenocarcinoma (READ, C-index = 0.606, *p* = 0.015), bladder urothelial carcinoma (BLCA, C-index = 0.581, *p* = 0.029), head and neck squamous cell carcinoma (HNSC, C-index = 0.543, *p* = 0.026), acute myeloid leukemia (LAML, C-index = 0.537, *p* = 0.026), colon adenocarcinoma (COAD, C-index = 0.570, *p* = 0.020), and cutaneous melanoma (SKCM, C-index = 0.550, *p* = 0.048). The mean C-index across all 30 cancers was 0.575. Across the entire pan-cancer population, quintile stratification by *η* yielded a Q1-vs-Q5 log-rank *p* = 6.9 × 10^−32^ (Fig. 1D).

**Table 1.** Survival stratification by *η* across cancer types. Only cancers with p < 0.05 are shown. Full results for all 30 evaluable cancers are in Supplementary Table 4 and Fig. S1.

| Cancer | Abbrev. | $n$ | C-index | AUC | $p$ |
| --- | --- | --- | --- | --- | --- |
| Glioblastoma | GBM | 76 | 0.845 | 0.906 | $1.8 \times 10^{-6}$ |
| Prostate adenoca. | PRAD | 91 | 0.800 | 0.876 | $8.9 \times 10^{-3}$ |
| Kidney clear cell | KIRC | 505 | 0.649 | 0.700 | $7.4 \times 10^{-5}$ |
| Uterine endometrial | UCEC | 80 | 0.634 | 0.681 | $9.0 \times 10^{-3}$ |
| Lung SCC | LUSC | 86 | 0.618 | 0.655 | $2.3 \times 10^{-2}$ |
| Rectal adenoca. | READ | 250 | 0.606 | 0.607 | $1.5 \times 10^{-2}$ |
| Bladder urothelial | BLCA | 291 | 0.581 | 0.570 | $2.9 \times 10^{-2}$ |
| Colon adenoca. | COAD | 505 | 0.570 | 0.588 | $2.0 \times 10^{-2}$ |
| Cutaneous melanoma | SKCM | 361 | 0.550 | 0.576 | $4.8 \times 10^{-2}$ |
| Head & neck SCC | HNSC | 267 | 0.543 | 0.506 | $2.6 \times 10^{-2}$ |
| AML | LAML | 400 | 0.537 | 0.517 | $2.6 \times 10^{-2}$ |

**Figure 3.**
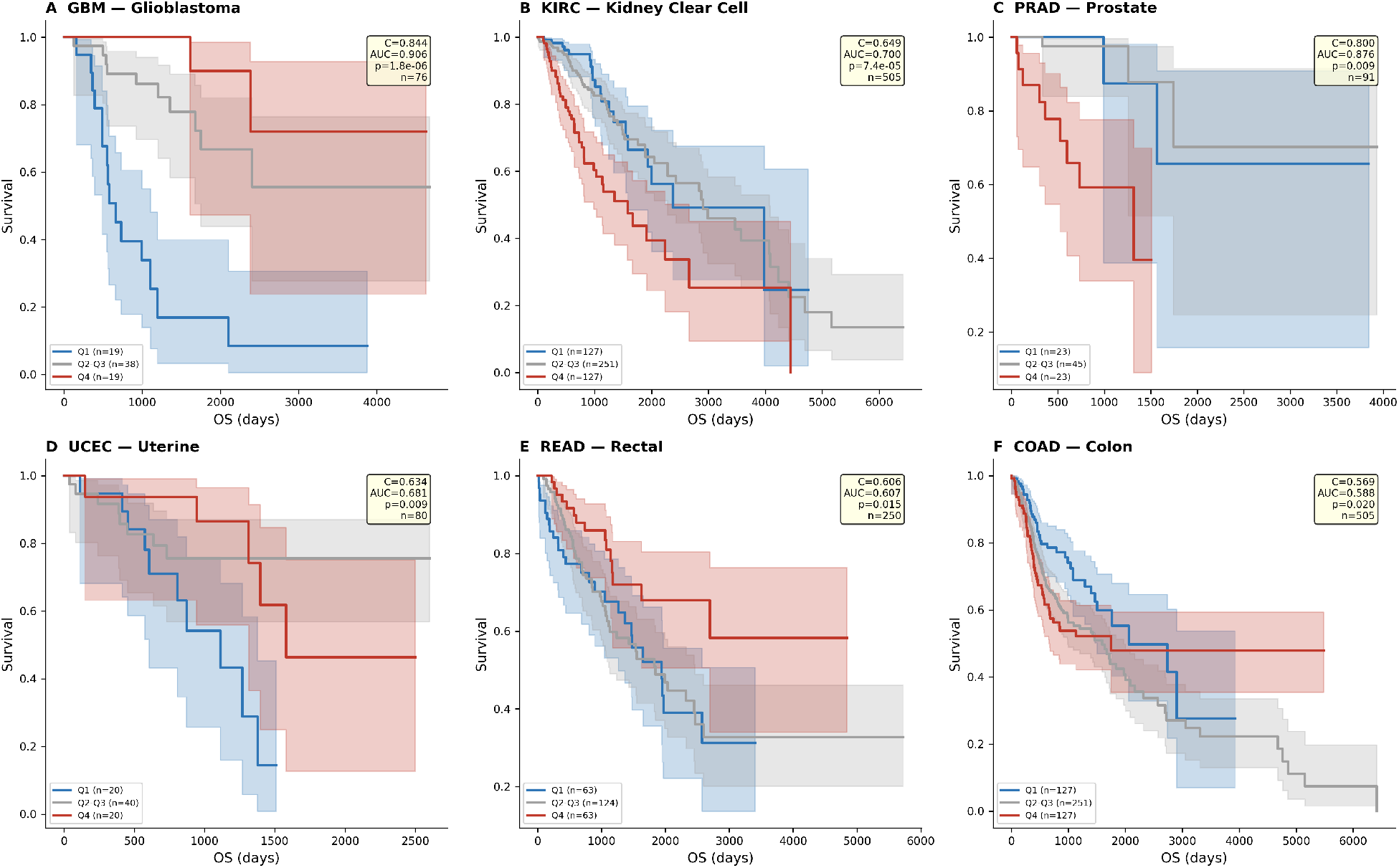
The spectral malignancy coordinate *η* stratifies overall survival across cancer types. Kaplan–Meier survival curves for the six cancer types with the strongest *η*–survival associations, stratified by *η* quartile: Q1 (lowest *η*, blue), Q2–Q3 (middle 50%, gray), and Q4 (highest *η*, red). Inset boxes report concordance index, time-dependent AUC, log-rank *p*-value (Q1 vs. Q4), and sample size. (A) Glioblastoma (C-index = 0.844, AUC = 0.906, *p* = 1.8 × 10^−6^, *n* = 76). (B) Kidney clear cell carcinoma (C-index = 0.649, AUC =0.700, *p* = 7.4 × 10^−5^, *n* = 505). (C) Prostate adenocarcinoma (C-index = 0.800, AUC = 0.876, *p* = 0.009, *n* = 91). (D) Uterine endometrial carcinoma (C-index = 0.634, AUC = 0.681, *p* = 0.009, *n* = 80). (E) Rectal adenocarcinoma (C-index = 0.606, AUC = 0.607, *p* = 0.015, *n* = 250). (F) Colon adenocarcinoma (C-index = 0.569, AUC = 0.588, *p* = 0.020, *n* = 505). Shaded bands indicate 95% confidence intervals.

The cancers in which *η* performed best are precisely those known to exhibit transcrip-tomic gradients along differentiation axes (Fig. 4): GBM (proneural–mesenchymal), KIRC (angiogenic–proliferative), UCEC (endometrioid–serous), and PRAD (luminal–basal). This is consistent with the biological identity of Mode 3 as a dedifferentiation axis: cancers whose clinical outcomes are driven by the degree of differentiation loss are the ones where the spectral coordinate is most informative.

**Figure 4.**
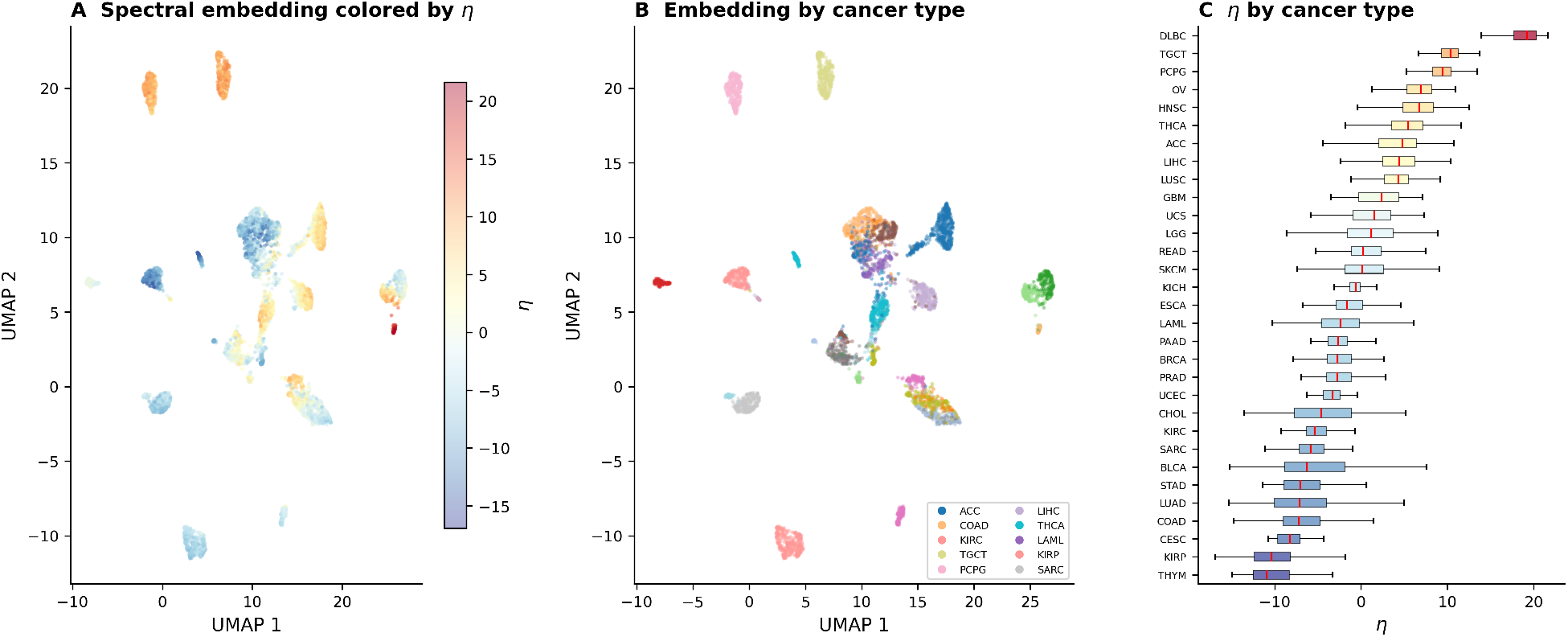
Spectral embedding and *η* distributions across cancer types. **(A)** UMAP embedding of 5,000 randomly sampled tumors projected onto the top 15 coordination modes, colored by the spectral malignancy coordinate *η*. The continuous gradient from blue (low *η*) to red (high *η*) crosses tissue-type boundaries, indicating that the coordinate captures a pan-cancer axis of coordination rather than tissue-of-origin identity alone. **(B)** Same embedding colored by cancer type, demonstrating that coordination modes recover tissuespecific clustering. Each island corresponds to a cancer type occupying a distinct region of the 15-dimensional coordination space. **(C)** Horizontal box plots of *η* by cancer type, sorted by median *η*. The wide range of medians across cancer types reflects the tissue-specific baseline of coordination architecture, while within-type spread reflects inter-patient heterogeneity along the spectral axis.

### External validation in non-overlapping TCGA samples

To assess generalizability, we projected the MLOmics-derived eigenvectors onto 2,480 non-overlapping TCGA Pan-Cancer Atlas samples (994 events) using 2,801 of 3,000 matched genes. No parameters were retrained; the scaler, eigenvectors, and mode selection were frozen from the discovery cohort. Across the full external cohort, quintile stratification by *η* maintained significant survival discrimination (Q1 vs. Q5 log-rank *p* = 8.5 × 10^−9^). At the individual cancer level, three of 12 evaluable types reached significance: breast carcinoma (C-index = 0.579, *p* = 0.024, *n* = 456), glioblastoma (C-index = 0.560, *p* = 0.046, *n* = 165), and lung squamous cell carcinoma (C-index = 0.545, *p* = 0.047, *n* = 193). Per-cancer C-indices were attenuated relative to the discovery cohort, consistent with cross-platform normalization differences and the loss of 199 unmatched genes. Notably, the pan-cancer coordination signal (*p* = 8.5 × 10^−9^) transferred robustly despite zero retraining, supporting the interpretation that the gene coordination structure captured by the Hamiltonian reflects biological organization rather than dataset-specific artifacts.

### The coordination threshold as a molecular point of no return

The monopolar architecture (*c* = 0.71) combined with the survival-associated secondary modes suggests a model of cancer progression as coordination collapse. In the early stages, a tumor’s transcriptome retains the multi-program architecture of its tissue of origin— differentiation programs, immune surveillance, and metabolic identity operate as independent coordination modes. As the tumor progresses, these programs become subordinated to the dominant coordination mode, and the secondary modes weaken. The spectral co-ordinate *η* measures a patient’s position along this trajectory: low |*η*| indicates retained multi-program coordination (the system is still in its distributed phase), while high |*η*| indicates that one or more secondary modes have been absorbed into the dominant program (the system has crossed the phase boundary into the ordered phase).

This interpretation is supported by the direction of the survival effect. In GBM (C-index = 0.845), patients with extreme *η* values—those farthest from the coordination centroid— have dramatically worse outcomes. The Q1-vs-Q4 Kaplan–Meier separation (Fig. 3A) shows that the bottom quartile of *η* achieves roughly 80% 4-year survival while the top quartile reaches less than 20%, consistent with the known proneural (favorable) to mesenchymal (unfavorable) gradient in glioblastoma biology.

The critical temperature *T* ^*^ provides a principled threshold for this transition. Below *T* ^*^, the Boltzmann distribution over coordination modes concentrates on the dominant program; above it, secondary programs become thermally accessible. A patient whose coordination profile places them below *T* ^*^ has, in physical terms, insufficient thermal energy to escape the dominant mode—their gene coordination has “frozen” into a single program. This is the molecular point of no return.

## Discussion

The central finding of this study is that the pan-cancer transcriptome, when analyzed through the lens of gene–gene coordination rather than gene–patient variance, reveals a strikingly simple architecture: one dominant coordination program captures 71% of all inter-gene coupling, with biologically interpretable secondary modes that predict survival across 11 cancer types. This simplicity is itself surprising. The pan-cancer landscape comprises 32 tissue types, thousands of driver mutations, and diverse microenvironmental compositions, yet the coordination structure reduces to a monopolar geometry with three informative orthogonal axes.

### Why coordination, not variance

The distinction between gene–gene coordination and sample–sample variance is the methodological core of this work, and it bears explicit discussion because it determines what the analysis can and cannot reveal.

Standard approaches—PCA, non-negative matrix factorization, consensus clustering— decompose the sample covariance matrix. Their eigenvectors are patient groupings: linear combinations of patients that maximize variance. This is the right tool for asking “which patients are different from each other?” but the wrong tool for asking “which genes move together?” A gene can have high variance across patients (because it is expressed in some tissue types and not others) without participating in any coordination program. Conversely, a gene with modest variance can be tightly coordinated with dozens of other genes within a functional module.

The Hamiltonian kernel *H*_*ij*_ measures a fundamentally different quantity: how similarly genes *i* and *j* behave across the patient population. Its eigenvectors are gene groupings— coordination programs—and its eigenvalues measure the tightness of that coordination. This is why the GSEA results are immediately interpretable: each eigenmode is a set of genes, and gene set enrichment directly queries what biological function that set encodes. By contrast, GSEA on PCA loadings typically yields diffuse, hard-to-interpret results because PCA loadings reflect variance contributions, not coordination structure.

### Biological interpretation of the coordination modes

The three survival-associated modes correspond to well-established axes of cancer biology, providing external validation of the framework:

Mode 3 (metabolic dedifferentiation) recapitulates the differentiation-state axis described in multiple cancer types. In renal cell carcinoma, this corresponds to the ClearCode34 angiogenic/proliferative gradient. In glioblastoma, it maps to the proneural–mesenchymal spectrum. The enrichment for estrogen response programs (NES = +2.19) on the differentiated pole suggests that hormone receptor signaling may serve as a broader marker of epithelial coordination integrity, even in non-hormone-driven cancers.

Mode 15 (immune polarization) separates adaptive cytotoxic immunity (IFN-*γ*, CD8B, CXCL9) from innate inflammatory programs (TNF-*α*/NF-*κ*B, IL-17). This axis is directly relevant to immunotherapy: tumors on the adaptive pole are candidates for checkpoint blockade, while those on the inflammatory pole may benefit from innate immune modulation. The enrichment for allograft rejection (NES = −2.37) on the adaptive pole is particularly notable, as this gene set is a validated surrogate for cytotoxic immune infiltration in the tumor microenvironment.

Mode 20 (tissue-identity dissolution) explains the strong performance of *η* in KIRC specifically. The renal transporters that dominate this mode’s gene loadings (SLC4A1, ATP6V0A4, CLCNKB) are functional markers of differentiated proximal tubule epithelium. Their coordinated loss indicates that the tumor has dissolved its tissue-specific program, which is known to be the primary driver of outcome in clear cell RCC.

### The phase transition interpretation

The statistical-mechanical framing of this analysis is not metaphorical. The Hamiltonian *H* is a real positive semi-definite operator with a well-defined eigenspectrum; the partition function, free energy, and heat capacity are computed from this spectrum by the standard formulas of statistical mechanics. The phase transition—the reorganization of the Boltzmann distribution from multi-mode to single-mode dominance—is a mathematical property of the eigenvalue distribution, not an analogy.

What is analogous is the mapping from physical to biological variables. In a magnetic system, the Hamiltonian encodes spin–spin interactions and the phase transition corresponds to spontaneous magnetization. In our framework, the Hamiltonian encodes gene–gene coordination and the “phase transition” corresponds to the collapse of multi-program coordination into single-mode dominance. The temperature *T* is a scanning parameter that reveals the structure of the eigenspectrum: at high *T*, all modes contribute equally; at low *T*, the dominant mode absorbs all weight. The critical temperature *T* ^*^ is the point at which this transition becomes sharp.

The biological claim is that cancer progression moves a tumor’s coordination landscape from the high-*T* (multi-program) regime toward the low-*T* (monopolar) regime, and that the spectral coordinate *η* measures position along this trajectory. This is testable in longitudinal data, where one would predict that serial biopsies of progressing tumors show increasing spectral concentration and decreasing secondary mode amplitudes.

### Limitations

Several limitations should be acknowledged. First, the analysis uses bulk mRNA expression, which conflates cell-intrinsic coordination with cell-composition effects. Single-cell resolution would distinguish whether the coordination modes reflect within-cell regulatory programs or between-cell-type compositional gradients. Second, although external validation in 2,480 non-overlapping TCGA samples confirmed the pan-cancer survival signal (*p* = 8.5 × 10^−9^), per-cancer C-indices were attenuated, with only 3 of 12 evaluable types reaching significance in the external cohort. This attenuation likely reflects cross-platform normalization differences and incomplete gene matching (2,801 of 3,000 genes), but overfitting to the discovery cohort cannot be excluded for cancer types with small sample sizes. Third, the Boltzmann-weighted version of *η* yielded near-uniform weights across the three survival modes (0.331, 0.335, 0.335), indicating that the temperature framework does not substantially differentiate the survival-associated modes from each other at the current spectral gap. The physics operates most powerfully at the macro level (the *c* = 0.71 monopolar finding and the existence of the phase boundary) rather than at the level of individual mode weighting. Fourth, 19 of 30 cancer types showed no significant survival association with *η* (Fig. S1). These may be cancers in which non-transcriptomic features (genomic alterations, epigenetic state, clinical stage) dominate prognosis, or cancers whose coordination axes are oriented differently from the pan-cancer modes.

### Implications

This work establishes gene–gene coordination analysis as a distinct analytical paradigm from the variance-based approaches that dominate current transcriptomic analysis. The coordination modes are directly interpretable as biological programs, their eigenvalues quantify the strength of inter-gene coupling, and the spectral concentration provides a single number that characterizes the organizational architecture of a cancer’s transcriptome. The spectral malignancy coordinate *η* offers a continuous, tissue-agnostic biomarker that measures not what genes are expressed but how they are organized relative to each other.

The immediate translational implication is that *η* could serve as a companion diagnostic for therapies that target specific coordination programs. A tumor on the dedifferentiation pole of Mode 3 might benefit from differentiation therapy; a tumor on the adaptive-immune pole of Mode 15 from checkpoint blockade; a tumor that has dissolved its tissue-identity coordination (Mode 20) from therapies that do not depend on tissue-specific targets. The spectral coordinate provides a principled basis for matching patients to mechanism-specific interventions.

## Materials and Methods

### Dataset

We used the MLOmics pan-cancer compendium, which comprises mRNA expression profiles for 8,314 tumors across 32 cancer types. Expression values were represented as normalized counts. We selected the 3,000 genes with highest variance across the full cohort. Overall survival data were obtained from the MLOmics clustering survival annotations (primary source) with backfill from the UCSC Xena Liu et al. (2018) unified survival endpoints, yielding 8,262 samples with complete overall survival data (2,286 events).

### Gene–gene coordination kernel (Hamiltonian)

Each gene’s expression profile was z-scored across all 8,314 patients, yielding a standardized profile g_*i*_ ∈ ℝ^8314^ for gene *i*. The gene–gene coordination kernel was computed as:

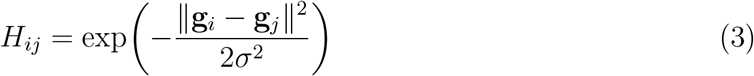

where *σ* was set to the median pairwise Euclidean distance between gene profiles (median heuristic; *σ* = 126.6). The resulting matrix *H* ∈ ℝ^3000×3000^ is symmetric and positive semi-definite, interpretable as the Hamiltonian of a system in which each gene is a degree of freedom and each entry *H*_*ij*_ measures the coordination strength between genes *i* and *j*. It was symmetrized as *H* ← (*H* + *H*^T^)*/*2 to correct for numerical asymmetry.

### Eigendecomposition

We computed the full eigendecomposition of *H* using `scipy.linalg.eigh,` retaining the top 100 eigenmodes (eigenvalues in descending order: *λ*_1_ ≥ *λ*_2_ ≥…). Each eigenvector v_*k*_ ∈ ℝ^3000^ is a coordination program—a weighting over genes that identifies a group of co-regulated transcripts. Its corresponding eigenvalue *λ*_*k*_ measures the coordination strength of that program.

#### Spectral parameters

##### Spectral concentration

*c* = *λ*_1_*/*∑ _*k*_ |*λ*_*k*_|, the fraction of total coordination captured by the dominant mode. Values near 1 indicate monopolar coordination; values near 1*/K* (for *K* retained modes) indicate uniform distribution.

##### Spectral gap

Δ*λ* = *λ*_1_ − *λ*_2_, measuring the separation between the dominant program and the strongest secondary program.

##### Critical temperature

*T* ^*^ = 1*/*Δ*λ*, defined by analogy with statistical mechanics as the temperature at which the Boltzmann distribution transitions from multi-mode to single-mode dominance.

##### Thermodynamic functions

The partition function 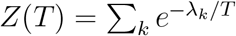, Helmholtz free energy *F* (*T*) = −*T* log *Z*(*T*), and heat capacity *C*_*v*_(*T*) =−*T ∂*^2^*F/∂T*^*2*^ were computed over the temperature range [0.01 *T* ^*^, 5 *T* ^*^] using the top 50 eigenvalues. The heat capacity peak identifies the temperature at which the eigenspectrum undergoes the sharpest redistribution of Boltzmann weights.

### Patient projection and spectral coordinate

Each patient’s z-scored expression profile x_*i*_ ∈ ℝ^3000^ was projected onto the coordination modes:

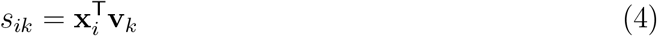

The score *s*_*ik*_ measures how strongly patient *i*’s expression profile aligns with coordination program *k*. The survival-associated modes were identified by screening the top 30 modes for concordance index (Harrell’s C-index) against overall survival across all patients with follow-up. For each mode, we computed both C-index(*T*, −*ψ*_*k*_, *E*) and C-index(*T*, +*ψ*_*k*_, *E*) and retained the maximum, to account for the arbitrary sign of eigenvectors. The top three modes by C-index were selected.

The spectral malignancy coordinate was defined as:

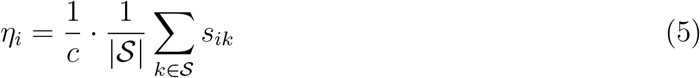

where 𝒮 = {3, 15, 20} is the set of selected modes. Division by spectral concentration *c* normalizes for the degree of monopolar dominance, analogous to dividing by a partition function.

### Survival analysis

For each of the 30 cancer types with ≥ 15 patients and ≥ 5 events, we evaluated *η* using:

1. **Concordance index**: Harrell’s C-index computed in both directions; the maximum was retained.
2. **Time-dependent AUC**: Binary outcome defined as event occurrence before the median event time; AUC computed and direction-corrected.
3. **Log-rank test**: Patients stratified into quartiles by *η*; Q1 vs. Q4 log-rank *p*-value.

### External validation

The MLOmics-derived eigenvectors were projected onto TCGA Pan-Cancer Atlas expression data obtained from the UCSC Xena TOIL RSEM pipeline. Of 3,000 genes in the eigenbasis, 2,801 matched by gene symbol. Samples overlapping with the MLOmics discovery cohort were excluded, yielding 2,480 independent samples with 994 events across 31 cancer types. Expression values were z-scored using the mean and standard deviation from the MLOmics discovery cohort (frozen scaler). The spectral coordinate *η* was computed identically to the discovery analysis with no parameter retraining.

### Gene set enrichment analysis

For each survival-associated coordination mode, genes were ranked by their eigenvector loading (the component of v_*k*_). Pre-ranked GSEA was run against MSigDB Hallmark 2020 and KEGG 2021 Human gene sets using `gseapy` with 1,000 permutations. Results were filtered at FDR *q <* 0.25.

### Software and reproducibility

All analyses were performed in Python 3.12 using NumPy, SciPy, scikit-learn, lifelines, statsmodels, and gseapy. Code and the processed alignment pickle (`pan_dataset_aligned.pkl`) are available at https://github.com/radres2019/hamiltonian-multiomics.

### Use of artificial intelligence

Large language model (LLM) assistance (Claude, Anthropic) was used in debugging analysis code and in typesetting this manuscript in LATEX. All scientific content, analytical decisions, and conclusions are the sole responsibility of the author.

#### Ethics statement

This study used only de-identified, publicly available transcriptomic and clinical datasets (MLOmics pan-cancer compendium and TCGA Pan-Cancer Atlas). No new human subjects research was conducted. Institutional review board approval was not required.

#### Funding

This work received no specific grant from any funding agency in the public, commercial, or not-for-profit sectors.

## Data Availability

Code is available at https://github.com/radres2019/hamiltonian-multiomics.. The MLOmics pan-cancer compendium is publicly available from Rappoport and Shamir (2018). Survival endpoints were obtained from the UCSC Xena platform (Liu et al., 2018). TCGA Pan-Cancer Atlas expression data were obtained from the TOIL Xena hub.

## Acknowledgments

We thank the creators and curators of the MLOmics pan-cancer compendium (Rappoport and Shamir, 2018) for making the transcriptomic data from 8,314 tumors across 32 cancer types publicly available. We also thank the UCSC Xena platform and the authors of Liu et al. (2018) for providing the unified survival endpoints derived from the TCGA Pan-Cancer Atlas clinical resource. Code is available at https://github.com/radres2019/hamiltonian-multiomics.

## Supplementary Material

**Figure S1.**
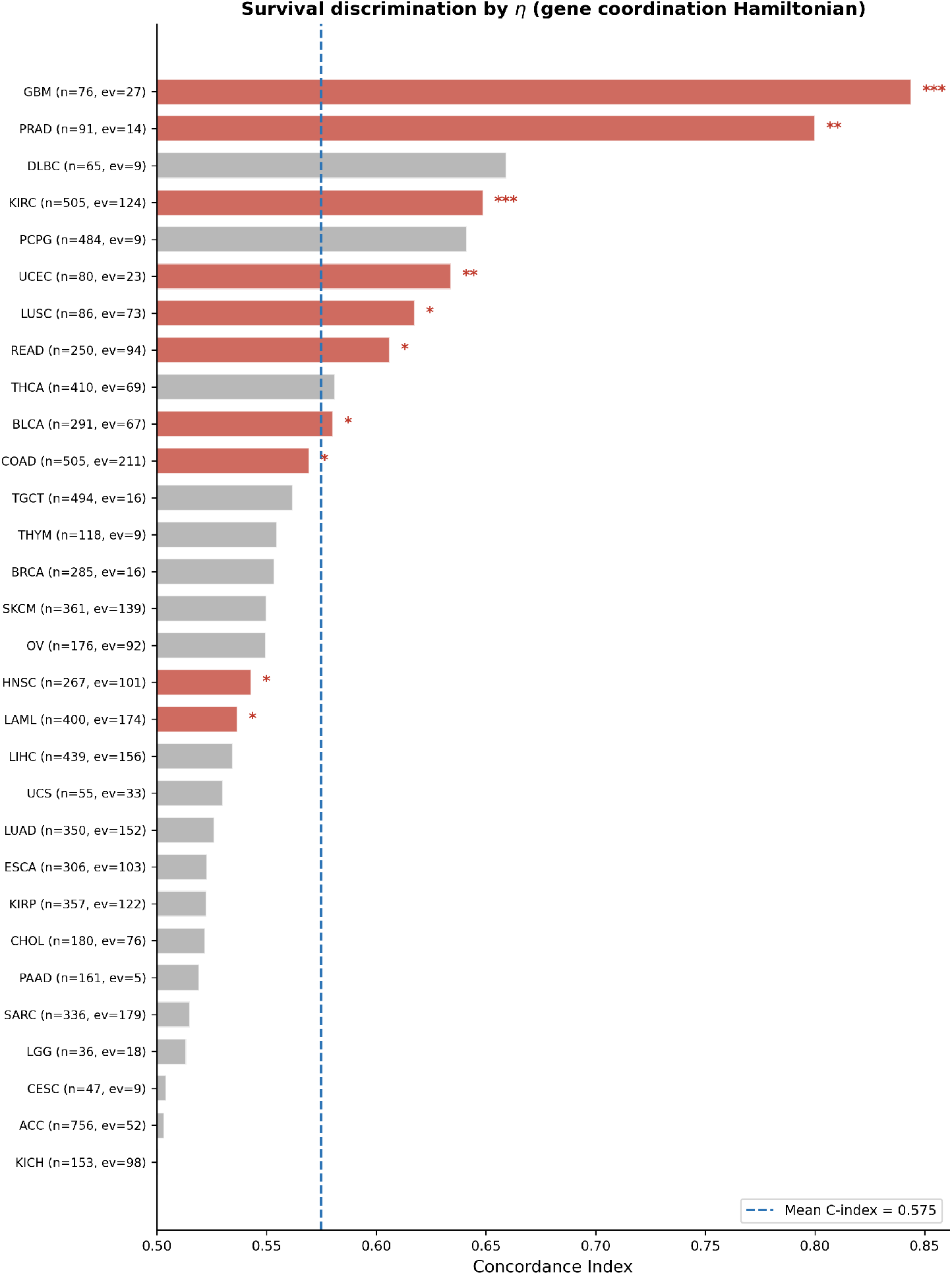
Survival discrimination by *η* across all 30 evaluable cancer types. Forest plot of concordance index for *η* in each cancer type. Red bars: significant (*p <* 0.05; 11 cancers); gray bars: not significant. Asterisks: ^*^*p <* 0.05, ^**^*p <* 0.01, ^***^*p <* 0.001. Dashed blue line: mean C-index = 0.575. Event counts (ev) shown alongside sample sizes (*n*).

